# Intraspecific interaction of host plant influences local distribution of specialist herbivores through metabolic alterations in leaves

**DOI:** 10.1101/2021.07.30.454541

**Authors:** Haruna Ohsaki, Atsuko Miyagi, Maki Kawai-Yamada, Akira Yamawo

## Abstract

1. Recent studies suggest that changes in leaf traits due to interactions between plants affect the resource utilisation and distribution of herbivores. However, this has not yet been confirmed experimentally. Here, we investigated the effects of phenotypic plasticity in leaf traits of *Rumex obtusifolius* (host plant) in response to the intra- and interspecific interaction on distribution of two leaf beetles, *Gastrophysa atrocyanea* (specialist herbivore) and *Galerucella grisescens* (generalist herbivore).
2. We investigated the local population density of *R. obtusifolius* plants and the presence of leaf beetles on the plants at five study sites. Leaf chemicals (condensed tannins and total phenolics) were compared between aggregated and solitary *R. obtusifolius* plants. To clarify the effects of the interaction environment of *R. obtusifolius* plants on their leaf traits and resource utilisation by leaf beetles, we conducted cultivation and preference experiments. Leaf chemicals (chlorophylls, organic acids, primary metabolites, condensed tannins and total phenolics) and preferences of adult leaf beetles were compared between intraspecific, interspecific plant interaction, or no-interaction treatments. Finally, we evaluated the effects of interaction between *R. obtusifolius* on leaf beetle distribution in mesocosm experiments.
3. In the field, the presence of the specialist leaf beetle, *G. atrocyanea*, was positively correlated with the local population density (rosette overlap ratio) of *R. obtusifolius* plants; however, no correlation was observed in the case of the generalist leaf beetle, *G. grisescens*. In the cultivation experiment, plants in the intraspecific interaction treatment increased their leaf contents of condensed tannins and total phenolics, and *G. atrocyanea* consumed more of these leaves than leaves in other treatments. Similar results were observed in the field. In the mesocosm experiment, larger numbers of *G. atrocyanea* were distributed on *R. obtusifolius* plants exposed to below-ground intraspecific interaction than on plants not exposed to intraspecific interaction.
4. Our results provide experimental evidence that leaf-trait changes in response to intraspecific interaction between host plants influence specialist herbivore distribution. This highlights the need to integrate plant–plant interactions into our understanding of plant–animal interactions.

## INTRODUCTION

To improve our understanding of plant–animal interactions, numerous ecologists have tried to predict herbivorous insect distribution by using the local population density of host plants. Root (1973) predicted that herbivores would be concentrated on host plants growing in high-density populations or monocultures (resource concentration hypothesis). This prediction has been supported by several studies (e.g., Stephens & Myers, 2012; Nerlekar, 2018). On the other hand, the possibility of an inverse distribution pattern, in which herbivores are concentrated on low-density or solitary host plants (Yamamura, 1999; Otway et al., 2005), has been proposed as the resource dilution hypothesis, and these predictions have also been supported by several studies (e.g., Fagundes et al., 2019; Coutinho et al., 2019). These conflicting patterns have been reported for several herbivore species, and some species have even been found to be unresponsive to resource distribution (Rhainds and English-Loeb, 2003; Tuller et al., 2013). Regardless, the mechanism that produces the uneven distribution of each herbivore species has not been elucidated.

Differences in the local population density of host plants are likely to reflect differences in the quality of the host plants, because the local population density of plants is linked to their interaction environments; host plants present at high density are exposed to direct intraspecific interaction. In contrast, host plants present at low density are exposed to direct interspecific interaction or no interaction. Many studies have reported that plant–plant direct interactions influence herbivory (Hambäck & Beckerman, 2003; Muiruri et al., 2019; Yamawo, 2021). For example, plant competition for resources induces plastic changes in the plants’ resource allocation; these changes can affect root or shoot growth. The changes in resource allocation can also influence the expression of leaf thickness, leaf mass per area, and primary (essential nutrients) and secondary (potentially plant-protective compounds) metabolites in the leaves (Bartron & Bowers, 2006; Broz et al., 2010; Mraja et al., 2011; Takigahira & Yamawo, 2019; Yamawo, 2021). Therefore, variations in the interaction environment induce changes in expression of the chemical traits of host plant leaves (Barton & Bowers, 2006; Broz et al., 2010; Mraja et al., 2011; Muiruri et al., 2019), so they are likely to influence leaf herbivory and the distribution of herbivores. In fact, several studies have strongly suggested that changes in leaf traits in response to differences in the interaction environment of host plants influence herbivory or herbivore distribution (e.g., Broz et al. 2010; Muiruri et al., 2019; Yamawo, 2021). However, to our knowledge, no experimental evidence has yet been provided for this effect.

Specialist herbivores are often attracted by secondary metabolites in their host plants; they adapt to these chemicals because they use them as cues to recognise the host plants (e.g., Wheat et al., 2007; Goodey et al., 2015). Brassicaceae plants produce glucosinolates to prevent herbivory by generalist herbivores; however, *Brevicoryne brassicae* (cabbage aphid), which specialises in Brassicaceae plants, prefers these glucosinolates (Titayavan & Altieri, 1990). Therefore, a high content of secondary metabolites induced in host plants by intraspecific interaction may attract the plants’ specialist herbivores. In contrast, generalist herbivores avoid secondary metabolites (e.g., alkaloids, phenolics and condensed tannins) in the leaves of host plants (e.g., Schoonhoven et al., 2005; Macel, 2011; Jeschke et al., 2017). Here, we hypothesised that intraspecific interaction increases the content of secondary metabolites in plant leaves, and that this increase would lead to the aggregation of specialist herbivores. In contrast, generalist herbivores may gravitate towards low-density host plants to avoid high contents of secondary metabolites. Therefore, differences in resource quality due to variations in local population density within a plant population could induce either a concentrated or a low-density distribution of herbivores, depending on the resource concentration (Root, 1973) or resource dilution (Otway et al., 2005) hypothesis, when compared with the distribution predicted on the basis of resource quantity alone (Fretwell & Lucas, 1969).

Here, we focused on *Rumex obtusifolius* L. (broad-leaved dock; Polygonaceae) as a host plant, and two leaf beetles, *Gastrophysa atrocyanea* Motschulsky (Chrysomelidae), which is a specialist herbivore of *Rumex* plants, and *Galerucella grisescens* (Joannis) (Chrysomelidae), which is a generalist herbivore of Polygonaceae plants (see details in Supplemental methods). To test our hypothesis, we investigated the relationships between the local population density of *R. obtusifolius* plants and the herbivores’ distributions in the field. Next, to clarify the effect of the interaction environment on leaf traits of *R. obtusifolius* plants and resource utilisation by the two leaf beetles, we conducted cultivation and preference experiments in adult leaf beetles. Finally, we evaluated the effects of intraspecific interaction of *R. obtusifolius* plants on the distribution of the leaf beetles by using a mesocosm experiment. On the basis of the results, we discuss the effects of plant–plant interaction on herbivore distributions.

## MATERIALS AND METHODS

### Field survey

To reveal the relationships between the local population density of *Rumex obtusifolius* and the distribution of leaf beetles, we conducted field surveys in April and May 2018, at a time when the populations of both leaf beetles are large. Five grasslands were selected as field-survey sites (5 April, Tomino-cho, Hirosaki City, Aomori Prefecture, 40°35ʹN 140°28ʹE; 13 April, Ozawa, Hirosaki City, Aomori Prefecture, 40°34ʹN 140°27ʹE; 28 April, Ohara, Hirosaki City, Aomori Prefecture, 40°34ʹN 140°26ʹE; 22 April, Nagoya City, Aichi Prefecture, 35°09ʹN 136°58ʹE; 5 May, Morioka City, Iwate Prefecture, 39°42ʹN 141°08ʹE, Figure S1). These sites are all at least 2 km apart. At each site we set up one square quadrat (Tomino-cho and Ohara, 10 × 10 m; Ozawa, 8 × 8 m; Iwate, 4 × 6 m; Nagoya; 4 × 4 m) including varying local population densities and sizes of *R. obtusifolius* plants. The maximum size of each quadrat was determined as 100 m^2^; in the case of small *R. obtusifolius* populations we adjusted the size of the quadrat downward to include all *R. obtusifolius* individuals.

#### Survey of local population densities of R. obtusifolius and herbivore distributions on R. obtusifolius

In each quadrat, a corner was used as the origin of two axes, x and y, which we used to plot coordinates. From the origin, we described the positions of all *R. obtusifolius* individuals, except for first-year seedlings that had cotyledons, to a precision of 1 cm by using a ruler. The longest rosette diameters of the described *R. obtusifolius* plants were recorded as the plant size. We also recorded the presence or absence of each herbivore species on each *R. obtusifolius* individual, regardless of the beetles’ developmental stages (egg, larva or adult).

By using these data, a bubble chart was created by converting the positions of plants into distributions on a map and the rosette sizes into bubble sizes (Figure S2). As an indicator of the local population density of *R. obtusifolius*, the area of one rosette overlapping with the rosettes of neighbouring individuals was calculated by using image analysis software (Adobe Photoshop Elements 2.0, Adobe Systems, San Jose, CA, USA). The overlap ratio (overlapping area/total rosette area) was used to represent the population density for analytical purposes. Because leaf beetles often retire into the soil around the host plants, making it difficult to evaluate their numbers accurately, we used binomial data (presence or absence) to analyse their distribution. The correlations between the overlap ratio of the rosettes and the presence of *Gastrophysa atrocyanea* or *Galerucella grisescens* were examined.

#### Measurement of leaf traits in field plants

To reveal the effects of local population density on the secondary chemicals of *R. obtusifolius* in the field, we measured the leaf secondary metabolites of *R. obtusifolius* plants that grew alone or were aggregated. In April 2018, leaves of *R. obtusifolius* plants were collected from three study sites in Aomori Prefecture, northern Japan (Hirosaki-city: 40° 35ʹN 140° 28ʹE, Fujisaki-city: 40° 39ʹ N 140° 29ʹ E, Itayanagi-city: 40° 40ʹ N 140°28ʹ E). Each site was at least 10 km apart from the next site. To exclude the effects of reproduction and leaf damage, we selected non-flowering individuals that had no herbivores and no leaf damage. An *R. obtusifolius* plant was defined as “Solitary” when there were no conspecific individuals within 30 cm from the edge of the widest rosette (*N* = 15), and *R. obtusifolius* plants with five or more conspecific individuals within a radius of 1 m from the centre of the plant were defined as “Aggregated” (*N* = 25). The widest rosettes of these plants were about 30 cm in diameter. We selected the youngest, fully expanded leaves. These leaves were analysed for secondary metabolites, namely the contents of total phenolics and condensed tannin, which are well known as major secondary metabolites in the *Rumex* genus (Feduraev et al., 2019). We measured the leaf contents of total phenolics and condensed tannins in accordance with the methods of Feeny (1970) and Dudt and Shure (1994).

#### Leaf beetle choice experiment using leaf sections from naturally growing plants

In April 2018, Solitary and Aggregated *R. obtusifolius* plants (85 individuals each) with rosette diameters of about 30 cm were selected at random in Hirosaki City. We collected the youngest fully expanded leaves from the plants. We cut one 2-cm piece from the base of each collected leaf. A wet filter paper (8 cm in diameter) was placed in a covered Petri dish (8.5 cm in diameter), and a piece of leaf from a Solitary plant and a piece of leaf from an Aggregated plant were placed on it with one adult of *G. atrocyanea* or *G. grisescens*. The Petri dishes were kept in a growth chamber (25 °C, 12L, 12D). After 24 h, the damage to each leaf piece was estimated by image analysis. More details of the methods are given in the Supplementary methods.

### Cultivation experiments

#### Cultivation design

To examine the effects of the interaction environment on leaf traits and leaf beetle preferences, we conducted cultivation experiments. To prepare enough samples to measure leaf traits and leaf beetle preferences, two experiments were conducted. Experiment 1 was conducted in 2017 to estimate the effects of interaction environment on leaf secondary metabolite contents and plant biomass. Experiment 2 was conducted in 2019 to estimate the effects of interaction environment on leaf primary metabolite and chlorophyll contents and leaf beetle preferences.

In September 2016, a total of more than 700 seeds of *R. obtusifolius* were collected from four individual plants in the field in Hirosaki City. Each individual was separated by at least 2 km. As interspecific competitors, we focused on *Plantago asiatica* L*., Trifolium repens* L. and *Festuca ovina* L. These species are the dominant competitors of *R. obtusifolius* in Japan (Ohsaki, 2020). A total of 100 seeds of *P. asiatica* were collected from two individuals in the field in Aomori Prefecture. A total of 100 seeds of *T. repens* were collected from individuals in the field in Saga Prefecture. For *F. ovina*, commercially available seeds (Kaneko Seeds Co., Gunma, Japan) were used. The seeds were stored in a refrigerator at 4 °C until the experiments began. Seeds from each mother plant were mixed and sown on the surface of wet sand (2 cm deep) during March 2017 for Experiment 1 and during March 2019 for Experiment 2. The containers were kept in a growth chamber (25 °C, 12L, 12D). All plants had developed their first true leaves by the beginning of the experiment.

In April 2017 and 2019, to obtain the focal plants, we planted one *R. obtusifolius* seedling in each pot (10.5 cm diameter × 9 cm high) containing seed-free garden soil (Mori Sangyo Co., Hokkaido, Japan). These pots were assigned to three interaction treatments: no-interaction treatment as a control (2017, *N* = 49; 2019, *N* = 35), intraspecific interaction treatment (2017, *N* = 66; 2019, *N* = 66) and interspecific interaction treatment (2017, *N* = 153; 2019, *N* = 98). In the no-interaction treatment, to provide a volume of soil similar to that used in the interaction treatment for each plant, the pots were divided into halves with a plastic plate to block any below-ground interaction, and one seedling of *R. obtusifolius* was planted in each half of the pot. In the intraspecific interaction treatment, we planted another *R. obtusifolius* seedling beside the focal plant as a competitor with no plastic plate. In the interspecific interaction treatment, a seedling of another species (*P. asiatica*, 2017, *N* = 58; 2019, *N* = 35; *T. repens,* 2017, *N* = 41; 2019, *N* = 30; *F. ovina*, 2017, *N* = 54; 2019, *N* = 33) was planted next to the target *R. obtusifolius* seedling. In the interaction treatment, the distance between seedlings was about 2 cm. All pots were placed randomly and maintained in the growth chambers (25 °C, 12L, 12D) and watered once a day for 30 days.

#### Measurement of leaf traits in cultivated plants

##### Experiment 1

After 30 days, we analysed total phenolics and condensed tannins. Plants were harvested and dried at 50 °C for 3 days. The plants were then weighed on an electronic balance to the nearest 0.1 mg. The leaves were used to analyse total phenolics and condensed tannins by using the methods in field survey.

##### Experiment 2

After 30 days, we measured chlorophyll content and five organic acids as plant nutrients (see details in Supplementary methods). The chlorophyll content reflects the plant’s nitrogen concentration and has been found to indirectly affect herbivore survival and distribution (Scheirs & De Bruyn, 2004, Sousa-Souto et al., 2018). Also, organic acids in the plant are necessary for the optimal development of phytophagous insects (Offor, 2010). Therefore, by measuring these, we examined changes in nutrient condition in response to interactions between plants.

#### Leaf beetle choice experiment

To reveal whether changes in leaf chemical contents induced in *R. obtusifolius* by interaction influenced the preferences of leaf beetles, we conducted choice experiments with the *R. obtusifolius* leaves used in cultivation experiment 2. The combinations of leaf pairs were as follows: intraspecific interaction versus interspecific interaction; interspecific interaction versus no-interaction treatment; and no-interaction treatment versus intraspecific interaction. The experimental design and conditions were similar to that described for the choice experiment using field leaves (see Supplementary methods).

### Mesocosm experiments

To determine the effects of the interaction environment of *R. obtusifolius* on the distribution of *G. atrocyanea*, we conducted a mesocosm experiment in November 2019 and July 2020. In November 2019, we estimated the effects of intraspecific interaction of *R. obtusifolius* plants on the distribution of leaf beetles by using plants of the same patch size in a “one-to-one-pot experiment” (Figure 1). In July 2020, we conducted a “one-to-three-pot experiment” to clarify the effects of intraspecific interaction of *R. obtusifolius* on the distribution of leaf beetles by adding the effects of patch size (i.e., resource amount) of the host plants.

**Figure 1.**
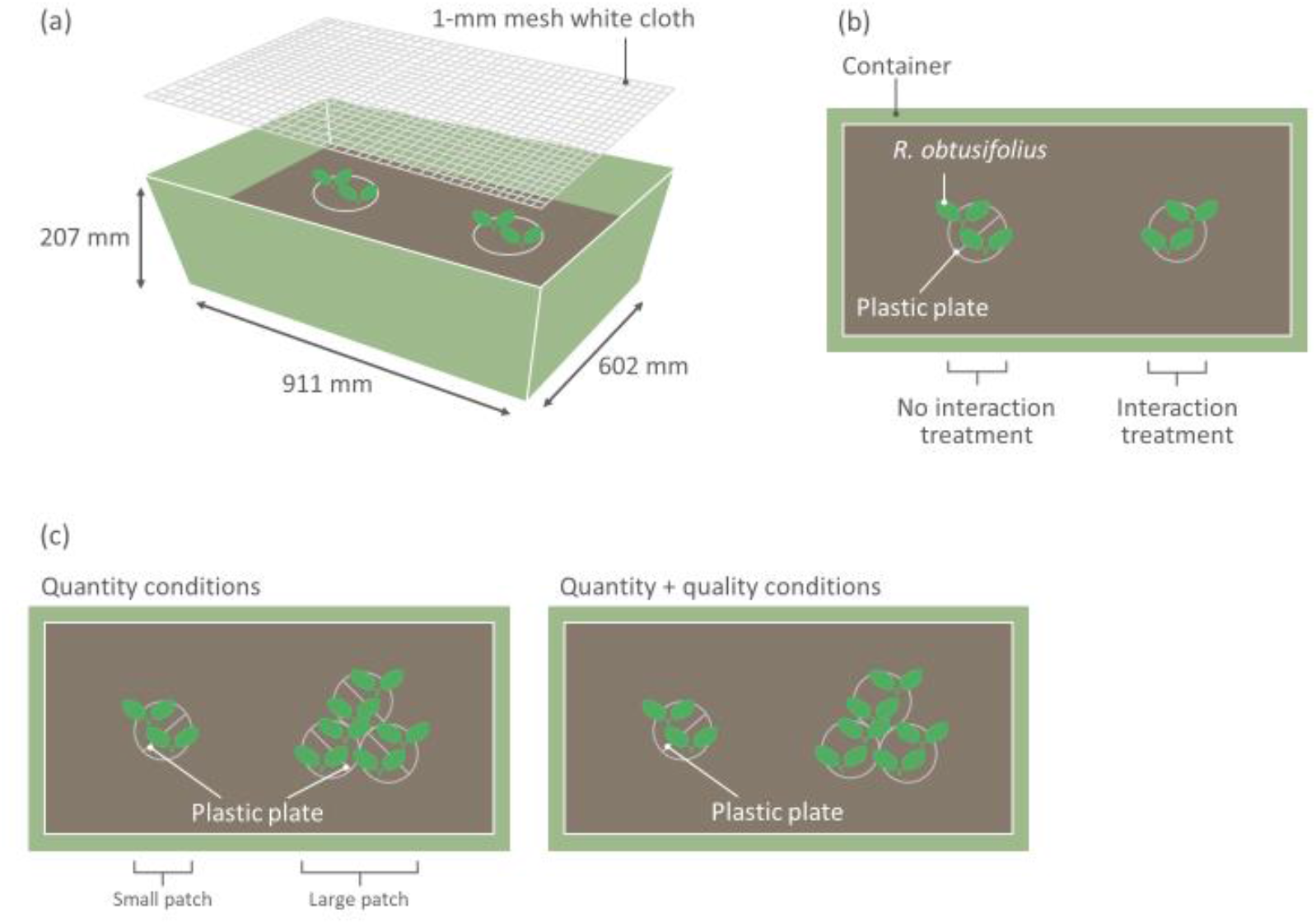
(a) Experimental setup in mesocosm experiment. In all containers, the area around the pots was filled with soil to a depth of 15 cm to allow the beetles free access to the plants, as in the field. (b) In the one-to-one-pot experiment, the interaction and no-interaction treatment pots were placed 30 cm away from each other in the container. (c) In the one-to-three-pot experiment, two sets of conditions were set up, namely “quantity conditions” and “quantity + quality conditions.” Under quantity conditions, two patch sizes were created by using four no-interaction pots. Under quantity + quality conditions, two patch sizes were created by using one no-interaction pot and three interaction pots. The distance between the large and small patches was 30 cm in each container.

We used two types of pot, namely interaction-treatment pots and no-interaction-treatment pots. In both types of treatment pot, two seedlings of *R. obtusifolius* were planted. The no-interaction-treatment pot was divided in half by a plastic plate to block below-ground interaction between *R. obtusifolius* plants. The interaction-treatment pot allowed below-ground interaction between *R. obtusifolius* plants, because this type of pot had no plastic plate (see Supplementary methods).

#### One-to-one-pot experiment

We prepared 20 containers (911 mm × 602 mm × 207 mm). In each container, an interaction-treatment pot and a no-interaction-treatment pot were placed 30 cm away from the edge of the pot. The containers were surrounded by soil to a depth of 15 cm to allow the beetles free access to the pods, as they would have in the field (Figure 1 a, b). For data analysis, each container was allocated an ID. Five *G. atrocyanea* females were released on the soil in the centre of each container, the top of which was then covered with 1-mm-mesh white cloth. The containers were placed in a greenhouse (15 °C), and the numbers of beetles on the plants were counted after 24 h.

#### One-to-three-pot experiment

In this experiment, we set up two types of conditions, namely “quantity conditions” (25 containers) and “quantity + quality conditions” (24 containers). For the quantity conditions, we set up patches of two sizes by using four no-interaction-treatment pots. We placed three pots together to represent large patches and one pot by itself to represent small patches (Figure 1c). For the quantity + quality conditions, we set up patches of two sizes by using one no-interaction-treatment pot and three interaction-treatment pots; the three interaction-treatment pots represented large patches and the single no-interaction-treatment pot represented small patches. In all containers, pots were set up as in the one-to-one-pot experiment and under the same controlled conditions. For the quantity conditions 125 beetles were used, and 120 beetles were used for the quantity + quality conditions.

### Statistical analysis

All statistical analyses were performed by using R v.3.6.1 software (R Development Core Team, 2019). All data met the statistical assumptions of normality and homoscedasticity according to the Kolmogorov–Smirnov test and F-test, and statistical analyses were performed appropriately depending on the data set structure. All tests were two tailed. The significance level was set at 0.05.

### Field survey data analysis

#### Survey of herbivore distribution on R. obtusifolius

We analysed the effects of the local population density of *R. obtusifolius* on the distribution of leaf beetles by using generalised linear mixed models (GLMMs) with a binomial distribution and logit function, followed by the Chi-square test. The models included presence or absence of leaf beetles as fixed terms and overlap ratio of *R. obtusifolius* rosettes for each plant, species of leaf beetles, and their interaction as explanatory variables. When the relationship between overlap ratio of rosette area and presence or absence of leaf beetles differed between leaf beetle species, the relationship between these was analysed for each beetle species. Site ID was included as a random effect in these models. False discovery rate (FDR) correction for multiple comparisons was then applied.

#### Measurement of leaf traits in field plants

Leaf chemical traits (content of condensed tannin or total phenolics) were compared between Solitary and Aggregate plants by using GLMMs with Gaussian distribution and an identity link, followed by an F-test; the models included leaf chemical traits as fixed terms and plant density (Solitary or Aggregated) as an explanatory variable. Site ID was included as a random effect in the models. FDR correction for multiple comparisons was then applied.

#### Leaf beetle choice experiment using leaf sections from naturally growing plants

Consumed areas of leaves were compared between local *R. obtusifolius* population densities for each leaf beetle species. We used GLMMs with Gaussian distribution and an identity link, followed by an F-test; the models included area consumed by leaf beetles as a fixed term and plant density (Solitary or Aggregated) as an explanatory variable. Petri dish ID was included as a random effect in the models. FDR correction for multiple comparisons was then applied.

### Cultivation experiments data analysis

#### Measurement of leaf traits in cultivated plants

We used Gamma distributions for the dry weights of plants, Gaussian distributions for the chlorophyll content of leaves, and Poisson distributions for the leaf contents of condensed tannin and total phenolics. We compared plant dry weights and leaf traits (condensed tannin, total phenolics and chlorophyll content) between the cultivation treatments by using GLMMs. Gamma or Poisson distributions with a log link followed by a Chi-square test were applied, and Gaussian distributions with an identifying link followed by an F-test were applied. These models included each plant trait as fixed terms and interaction treatment (no-, intraspecific or interspecific interaction treatment) as an explanatory variable. Parent plant ID was included as a random effect in the models. When there was an interaction effect between each plant trait and interaction treatment, we conducted multiple comparisons by FDR correction.

Organic acids were analysed by using a principal component analysis (PCA) based on the correlation matrix of variables. Scores on the first (PC1) and second (PC2) axes of the PCA were compared between interaction treatments by using GLMMs with a Gaussian distribution and an identity link, followed by an F-test. The models included PC1 or PC2 as fixed terms, interaction treatment (no-, intraspecific or interspecific interaction treatment) as an explanatory variable and parent plant ID as a random effect. When there was an interaction between PC1 or PC2 and interaction treatments, we conducted multiple comparison by using FDR correction.

#### Leaf beetle choice experiment

The leaf area consumed by the leaf beetles was compared between interaction treatments (no-, intraspecific or interspecific interaction). Data sets for female beetles were analysed by using GLMMs with a Gamma distribution and a log link, followed by a Chi-square test, and data sets of male beetles were analysed by using the Wilcoxon signed-rank test because the data sets contained some 0 values. The analysis was conducted for each species of leaf beetle and for each sex of each species. FDR correction for multiple comparisons was then applied to each data set.

### Mesocosm experiments data analysis

#### One-to-one-pot experiment

The numbers of leaf beetles per patch were compared between cultivation treatments by using GLMMs with Gaussian distributions and an identity link, followed by an F-test; the models included number of leaf beetles on the patch as a fixed term and interaction treatment (interaction or no-interaction) as an explanatory variable. Container ID was included as a random effect in the models.

#### One-to-three-pot experiment

Number of leaf beetles per patch or number of leaf beetles per pot (representing leaf beetle density) was compared between patch sizes (quantity) and cultivation conditions (quality) by using GLMMs with Poisson distributions and a log-link, followed by a Chi-square test; the models included number of leaf beetles per patch or per pot as fixed terms and patch size (small or large), cultivation conditions (quantity or quantity + quality condition) and their interaction as an explanatory variable. Container ID was included as a random effect in these models. When there was an interaction between patch size and cultivation conditions, we conducted multiple comparisons by FDR correction.

## RESULTS

### Field survey

More than 60 *R. obtusifolius* individuals were growing within each quadrat; the major herbivores were *G. atrocyanea* and *G. grisescens* (Table 1). The relationship between the overlap ratio of *R. obtusifolius* rosettes and the presence of leaf beetles differed among leaf beetle species (*x^2^* = 81.032, *df* = 2, *P* < 0.001). There was a significant positive correlation between the overlap ratio of *R. obtusifolius* rosettes and the presence of *G. atrocyanea* (estimate coefficient = 0.786, *x^2^* = 12.764, *df* = 1, *P* < 0.001). In contrast, the presence of *G. grisescens* was not significantly correlated with the overlap ratio of *R. obtusifolius* rosettes (estimate coefficient = –0.456, *x^2^* = 2.451, *df* = 1, *P* = 0.117).

**Table 1.**
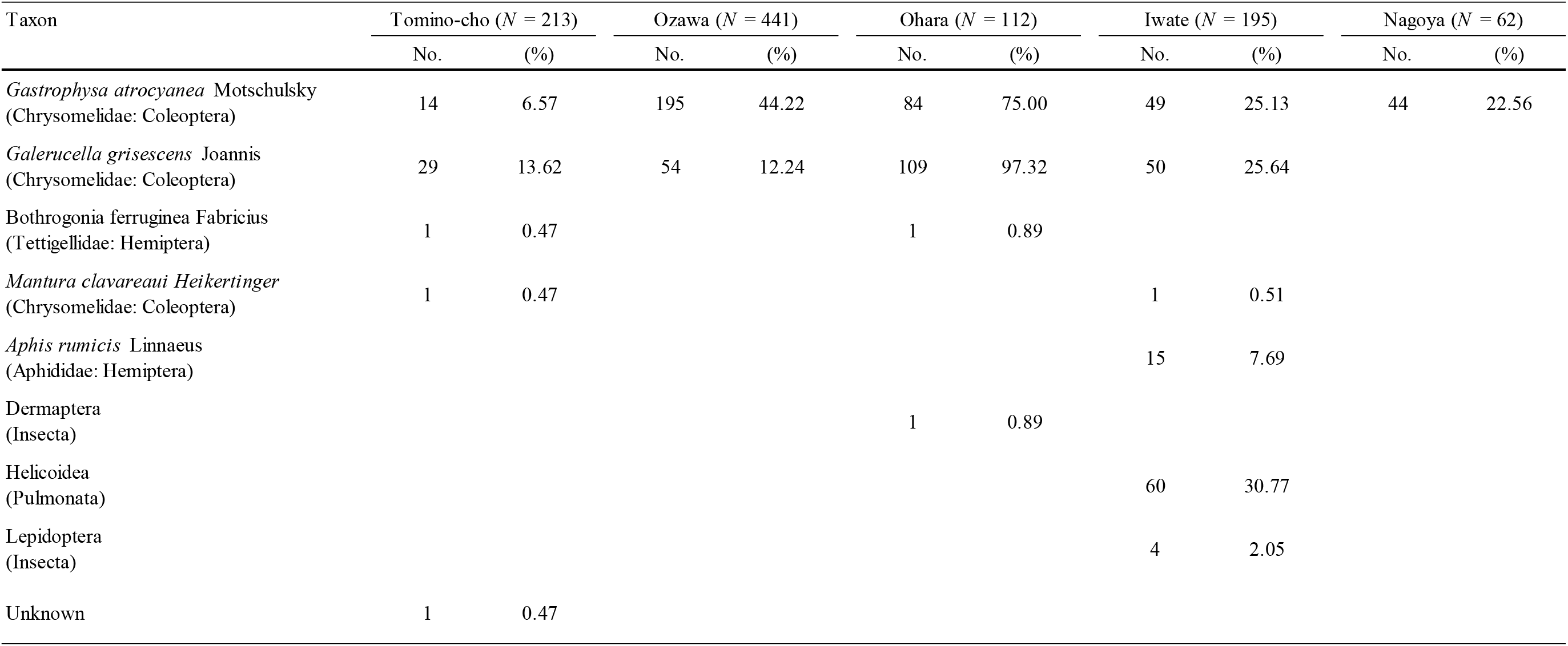
Herbivores of *Rumex obtusifolius* and numbers and proportions of infested plants.

Contents of total phenolics and condensed tannin tended to be higher in Aggregated plants than in Solitary plants, but this relationship did not reach statistical significance (total phenolics, *F* = 3.910, *P* = 0.096, Figure 2a; condensed tannin, *F* =4.882, *P* = 0.067, Figure 2b). Females of *G. atrocyanea* consumed significantly more leaf tissue from Aggregated plants than from Solitary plants (*F* = 7.837, *P* = 0.037, Figure 3a). In contrast, for males of *G. atrocyanea* (*F* = 1.779, *P* = 0.323, Figure 3b) and for both sexes of *G. grisescens* (female, *F* = 1.421, *P* = 0.323, Figure 3c; male, *F* = 0.494, *P* = 0.490, Figure 3d), there were no differences in the area of feeding damage between Aggregated and Solitary leaves.

**Figure 2.**
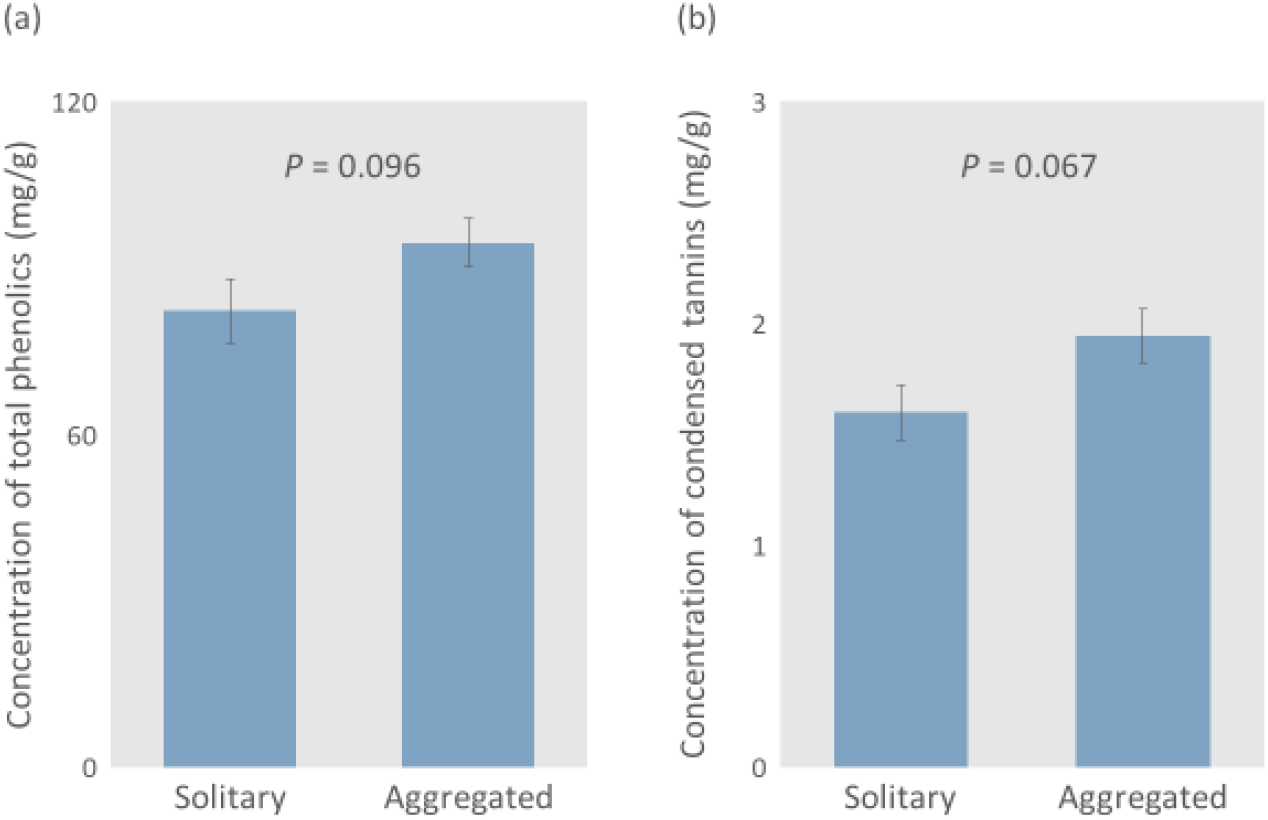
Contents of (a) total phenolics and (b) condensed tannin in Solitary (*N* = 15) and Aggregated (*N* = 25) plants. Bars represent SE. *P*-values are for the results of GLMM analysis.

**Figure 3.**
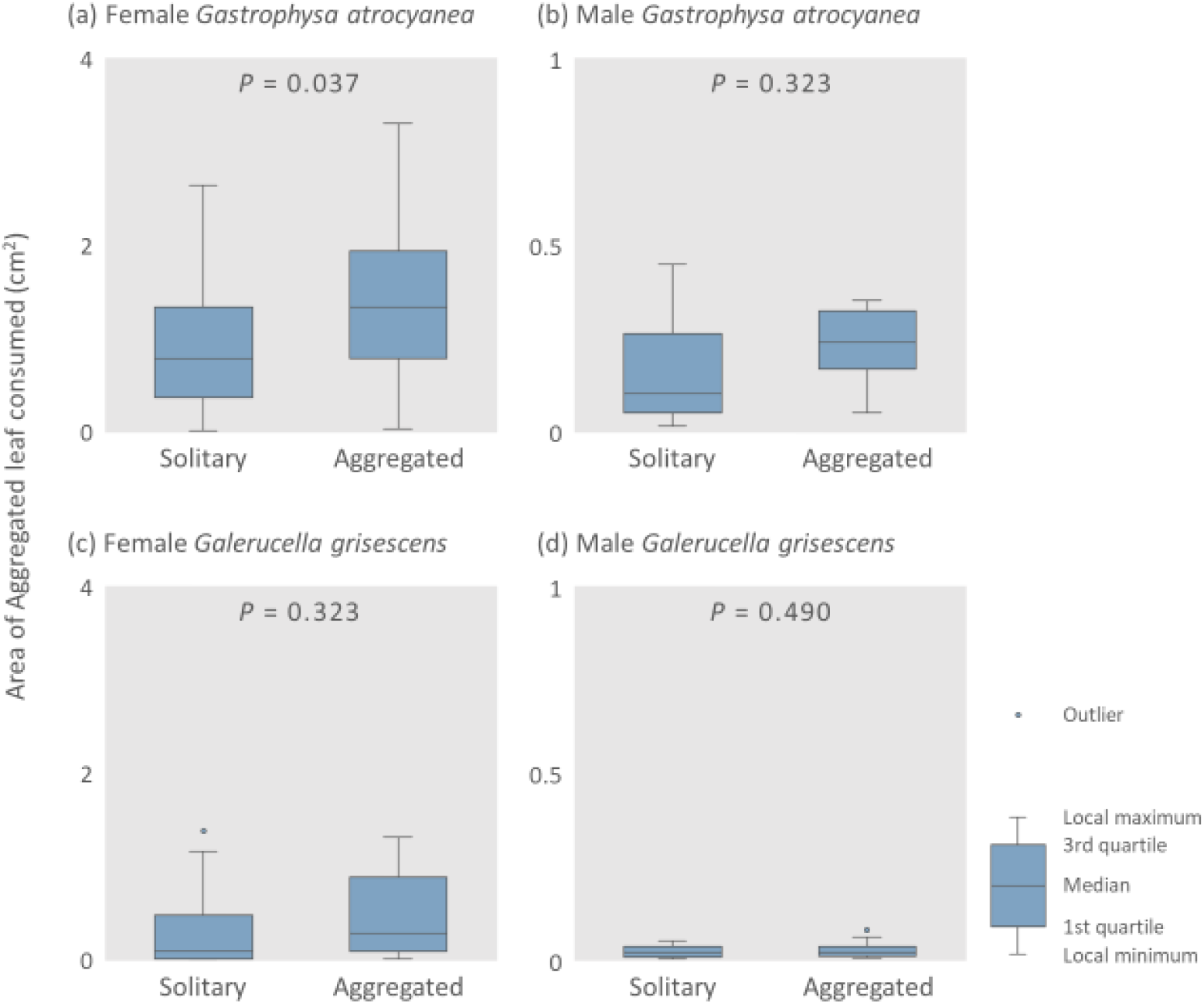
Areas of leaf consumed by (a) female and (b) male *Gastrophysa atrocyanea* and (c) female and (d) male *Galerucella grisescens* in the choice experiment using leaves from the field. *P*-values are for the results of GLMM analysis.

### Cultivation experiments

The biomass and chlorophyll content of *R. obtusifolius* did not differ among interaction treatments (biomass: *x^2^* = 7.081, *df* = 4, *P* = 0.132, Figure 4a; chlorophyll: *F* = 1.444, *P* = 0.239, Figure 4b). The contents of total phenolics of *R. obtusifolius* differed significantly among treatments; they were higher in the order of intraspecific, no-, and interspecific interaction treatment (Figure 4c). Plants subjected to the intraspecific interaction treatment had a significantly higher content of condensed tannins than those undergoing the no- or interspecific interaction treatments (Figure 4d).

**Figure 4.**
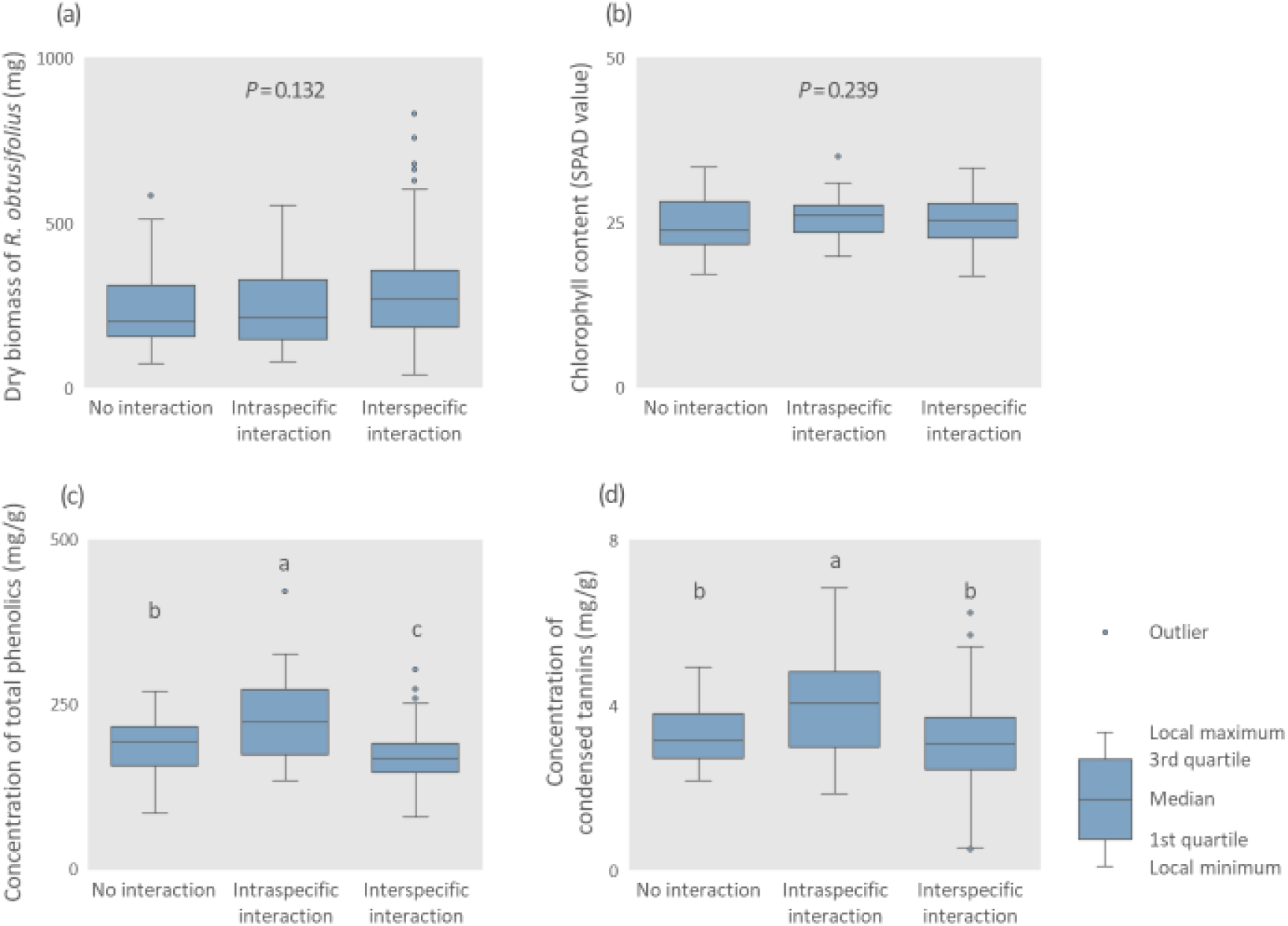
Dry biomass and leaf traits of *Rumex obtusifolius* in the cultivation experiment. (a) Dry biomass of whole plant, (b) chlorophyll content, (c) content of total phenolics and (d) content of condensed tannins in leaves. Different letters denote significant differences (GLMM, *P* < 0.05).

We found that PC1 and PC2 explained 61.0% and 20.7%, respectively, of the total variance of the organic acid composition data. The PC1 value did not differ among interaction treatments (*F* = 2.068, *P* = 0.154, Figure S3a). Plants subjected to the no-interaction treatment had significantly lower PC2 values than those undergoing the intraspecific or interspecific interaction treatments (Figure S3b).

Females of *G. atrocyanea* consumed significantly more leaf tissue from the intraspecific interaction treatment plants than from the no-interaction plants (*x^2^* = 5.470, *df* = 1, *P* = 0.029, Figure 5a) and from the interspecific interaction plants (*x^2^* = 6.064, *df* = 1, *P* = 0.029, Figure 5b). There was no significant difference between the no interaction and interspecific interaction treatments in terms of the area of leaf eaten by females of *G. atrocyanea* (*x^2^* = 1.832, *df* = 1, *P* = 0.176, Figure 5c). For males of *G. atrocyanea*, there were no differences between treatments in the area of leaf consumed (no-interaction versus intraspecific interaction, *z* = 0.329, *P* = 1, Figure 5d; interspecific interaction versus intraspecific interaction, *z* = 2.139, *P* = 1, Figure 5e; no-interaction versus interspecific interaction, *z* = 0, *P* = 0.097, Figure 5f).

**Figure 5.**
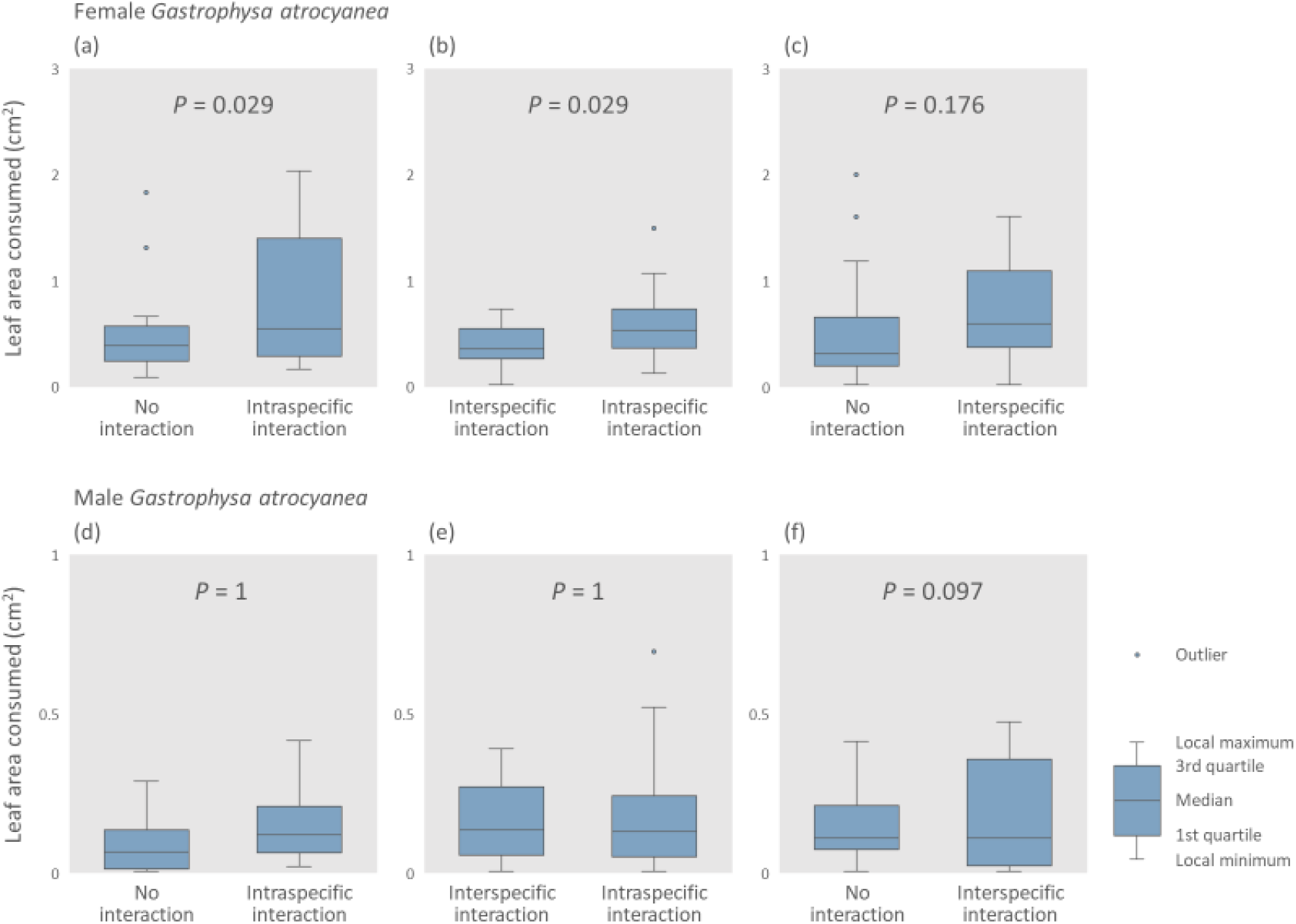
Leaf areas consumed by (a to c) female and (d to f) male *Gastrophysa atrocyanea* in the choice experiment using cultivated plant leaves. The combinations of leaf pairs of treatments were as follows: (a, d) no interaction versus intraspecific interaction; (b, e) interspecific interaction versus intraspecific interaction; (c, f) no interaction versus interspecific interaction.

### Mesocosm experiments

In the one-to-one-pot experiment, a significantly greater number of *G. atrocyanea* were distributed on the *R. obtusifolius* plants in the interaction treatment than in the no-interaction treatment (*F* = 5.556, *P* = 0.030, Figure 6a). In the one-to-three-pot experiment, the effect of patch size on the distribution of *G. atrocyanea* differed significantly with the cultivation conditions (*x^2^* = 6.540, *df* = 2, *P* = 0.038, Figure 6b). Under both types of cultivation condition, large patches had significantly more beetles than small patches (quantity conditions, *x^2^* =18.301, *df* = 1, *P* < 0.001; quantity + quality conditions, *x^2^* =55.474, *df* = 1, *P* < 0.001, Figure 6b). This trend was more pronounced under quantity + quality conditions. Moreover, the effect of patch size on the number of *G. atrocyanea* per pot (i.e., the leaf beetle density) differed significantly between cultivation conditions (interaction treatment × patch size; *z* = –2.067, *P* = 0.039, Figure 6c). Although the densities of leaf beetles in small and large patches were similar under quantity conditions (*z* = –0.308, *P* = 0.758), under quantity + quality conditions the large patches had a greater density of leaf beetles than small patches (*z* = 2.118, *P* = 0.034; Figure 6c).

**Figure 6.**
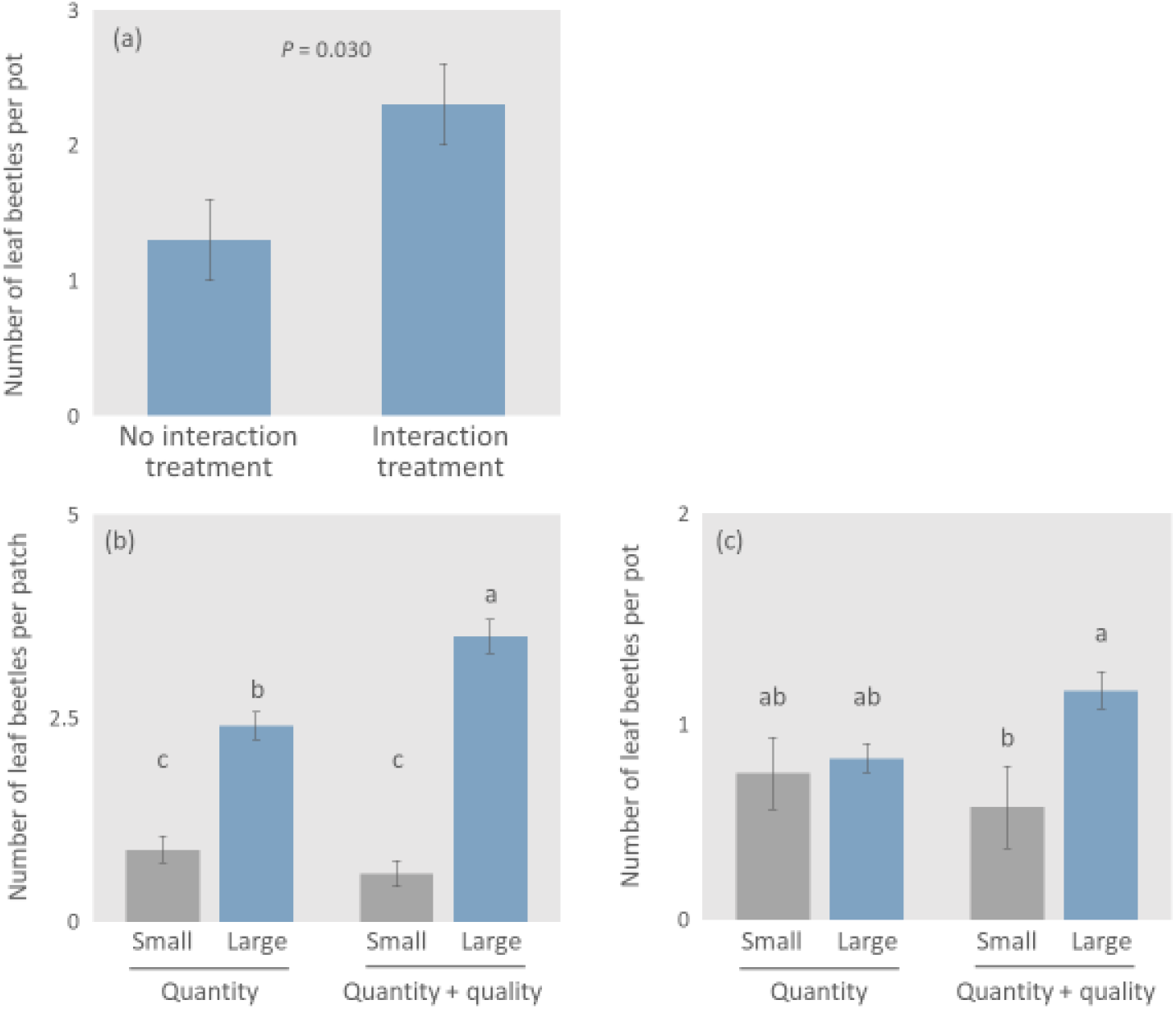
Numbers of leaf beetles in the mesocosm experiment. (a) Number of leaf beetles per pot in each treatment (no interaction or interaction treatment) in the one-to-one-pot experiment. (b) Number of leaf beetles per patch and (c) number of leaf beetles per pot in each patch (small and large patches) under each set of conditions (quantity and quantity + quality conditions) in the one-to-three-pot experiment. Bars represent SE. Different letters indicate significant differences (GLMM, *P* < 0.05).

## DISCUSSION

We found here that intraspecific interaction induced changes in the leaf metabolite contents of *Rumex obtusifolius* and affected resource utilisation by the specialist leaf beetle, *G. atrocyanea*, but not by the generalist leaf beetle, *G. grisescens*. In addition, we showed experimentally that this type of resource utilisation affected the distribution of *G. atrocyanea.* These results support our hypothesis, providing experimental evidence that differences in the local population density of the host plant led to plastic changes in leaf metabolite contents, affecting the resource utilisation and distribution patterns of specialist herbivores.

### Variations in leaf traits

In the field, Aggregated *R. obtusifolius* plants tended to have higher contents of total phenolics and condensed tannin than Solitary plants (Figure 2). This result suggests that aggregation of *R. obtusifolius* plants induced changes in leaf chemical traits. In fact, in the cultivation experiments, the contents of total phenolics and condensed tannin in the leaves of *R. obtusifolius* were significantly higher under intraspecific interaction conditions than under interspecific interaction or no-interaction conditions (Figure 4c, d). Increased contents of secondary metabolites in the presence of a conspecific neighbour have been reported in several plant species, and it has been suggested that metabolic alterations in leaves in response to intraspecific interaction are common in plants (Barton & Bowers, 2006; Ormeño et al., 2007; Broz et al., 2010, but see Kigathi et al., 2013). In many plant species intraspecific competition is more intense than interspecific competition (Adler et al., 2018), and such intraspecific competition causes limitation of soil nutrients and water (Craine & Dybzinski, 2013; Takigahira & Yamawo, 2019). It is well known that limitation of soil nutrients and water for plants induces the accumulation of secondary metabolites in the leaves (reviewed in Akula & Ravishankar, 2011). Thus, aggregation of *R. obtusifolius* plants may increase the leaf contents of secondary metabolites, such as total phenolics and condensed tannin, through soil resource competition.

Another possible hypothesis is that Aggregated plants invest more in defence than do Solitary plants through recognition of con-specific neighbours, because aggregated plants often consumed by specialist leaf beetles. Leaf-trait alteration based on neighbour recognition has also been reported in several plant species (Yamawo & Mukai 2020; Yamawo, 2015, 2021). Neighbour recognition can therefore be a cause of leaf-trait alteration in *R. obtusifolius* plants. However, the history of interaction between specialist leaf beetles and *R. obtusifolius* plants is weak, because *R. obtusifolius* is an exotic species in Japan. To understand the adaptive importance of leaf-trait alteration in *R. obtusifolius* plants, we would need to perform an additional study in a region to which *R. obtusifolius* is native.

The contents of primary metabolites are strongly affected by light conditions (Kitazaki et al., 2018). For example, experiments with lettuce, *Lactuca sativa*, have shown that the pattern of accumulation of primary metabolites, such as sugars and amino acids, is affected by light quality, intensity and exposure time (Kitazaki et al., 2018). In fact, our cultivation experiment found differences in the content of primary metabolites between no-interaction and interaction treatments (Figure S3b). Changes in the contents of primary metabolites in leaves may depends on presence of neighbour plants, regardless of the identity of neighbour.

### Preferences and distribution of leaf beetles

The local *R. obtusifolius* population density affected the amounts of leaf consumed by the specialist leaf beetle, *G. atrocyanea*. In the experiment using leaves from the field, females of *G. atrocyanea* preferred to consume the leaves of aggregated *R. obtusifolius* plants than of Solitary plants, despite similar quantities of leaves being provided for the beetles (Figure 3a). In the experiment using the leaves of cultivated plants, females of *G. atrocyanea* also preferred the leaves of *R*. *obtusifoliu*s plants exposed to intraspecific interaction over those of plants exposed to no interaction (Figure 5a). These preference pattern are consistent with the increases in the leaf contents of secondary metabolites (total phenolics and condensed tannins) (Figures 2 and 4c, d) but not with the variations in primary metabolites (Figures S3). Therefore, we concluded that females of *G. atrocyanea* selected leaves on the basis of increases induced in the leaf secondary metabolite content by the host plant’s interactive environment. *G. atrocyanea* beetles are specialist herbivores of *Rumex* plants (Suzuki, 1985). Many herbivore specialists use host-specific secondary metabolites for host searching or detecting (Schoonhoven et al., 2005; Ômura, 2018). This type of host searching may reflect the feeding preferences of *atrocyanea*. Females of *G. atrocyanea* lay eggs on the plants on which they feed, and the hatched larvae feed on the same plants. The larvae of *G. atrocyanea* require large amounts of food, and plants are often completely consumed (Suzuki, 1985). For this reason, the selection of aggregated plant leaves by *G. atrocyanea* females during the reproductive season is linked to the securing of food resources for the next generation. This may be associated with niche specialisation in coevolution among host plants and specialist herbivores (Schoonhoven et al. 2005; Abrahamson, 2008). In contrast, no preference was observed among males of *G. atrocyanea*, possibly because males use fewer resources than females with egg masses.

Do the preferences of leaf beetles affect the beetles’ distribution? Our mesocosm experiment provided robust evidence that changes in leaf traits based on intraspecific interaction can affect the distribution of the specialist leaf beetles, *G. atrocyanea* (Figure 6). When the plant patch sizes were similar, approximately 1.7 times more leaf beetles were distributed in the interaction treatment patch than in the no-interaction treatment patch (Figure 6a). Effects of interaction between host plants were also found in the one-to-three-pot experiment. Greater numbers of leaf beetles were distributed on the large patches than on the small patches, and this trend was more pronounced under quantity + quality conditions than under quantity conditions (Figure 6b). This finding is consistent with the distribution of *G. atrocyanea* in the field (Suzuki, 1985).

Moreover, in the conditions under which both the patch size and the competitive environment (and thereby leaf traits) differed, the leaf beetle density was significantly higher on large patches than on small ones, but it did not differ in conditions under which only the patch size differed (Figure 6c). Root (1973) predicted that “Herbivores are more densely distributed as the patch size of the feeding site increases (resource concentration hypothesis).” According to this prediction, the response of *G. atrocyanea* to differences in leaf traits leads to the concentration of beetles relative to the food resource. In other words, in this system, our results strongly suggested that differences in metabolic alterations in leaves through intraspecific interactions in plants induced the concentrated distribution of herbivores on resources.

In contrast, numbers of the generalist leaf beetle, *G. grisescens*, were not correlated with the local population density of *R. obtusifolius* in the field, and these beetles did not select the leaves of *R. obtusifolius* on the basis of the interaction environment. These results did not support our hypothesis that generalist herbivores accumulate on low-density host plants to avoid high levels of secondary metabolites. Generalist herbivores respond to a variety of chemicals besides those measured as leaf traits in this study (Schoonhoven et al., 2005; War et al., 2012). Perhaps other leaf traits, which could not be measured here, may have been involved in the preferences of *G. grisescens* and varied according to the interaction environment, thus masking the avoidance effect of secondary metabolites on the generalist leaf beetles. Another possible reason why the findings did not support our hypothesis is the effects of resource competition among herbivores. In some cases, resource competition among herbivores influences herbivore distribution (e.g., Suzuki 1986; Schoonhoven et al., 2005; Godinho et al., 2020). A previous study pointed out that *G. grisescens* is vulnerable to resource competition from *G. atrocyanea* (Suzuki, 1986). It may therefore prioritise the avoidance of competitors over plant availability when deciding where to feed (Suzuki, 1985). Several studies, as well as the resource dilution hypothesis proposed by Otway et al. (2005), have pointed out that herbivore density per plant may be higher when the host density is low (e.g. Yamamura, 1999). Our results suggest that these phenomena may be caused not only by differences in the local population density of the host plants but also indirectly by interactions with other herbivorous insects. To determine whether these results are general or specific to certain herbivores, several species, including generalists, may need to be tested.

## CONCLUSIONS

Our findings provide experimental evidence that intraspecific interaction between host plants affects specialist herbivore distribution. Many researchers have worked to unravel the relationship between the distribution of herbivores and the local population density of host plants. Some herbivores have shown a positive response to resource abundance, as in the resource concentration hypothesis proposed by Root (1973), whereas others, as in the resource dilution hypothesis proposed by Otway et al. (2005), have shown a negative response. These studies have focused on the amount of food available and have assumed that leaf traits are always constant. Our results indicate that herbivore responses to resource quantity and quality may interact with each other as factors governing herbivore distribution. Therefore, herbivore responses to the local population density of host plants can be understood from a plant–plant interaction perspective, highlighting the need to integrate plant–plant interactions into our understanding of plant–animal interactions in nature.

## ACKNOWLEDGEMENTS

This work was supported by JSPS Grants-in-Aid for Scientists (grant no. 18K19353 and 19H03295 to AY) from the Japan Society for the Promotion of Science and a Sasakawa Scientific Research Grant (Study Number: 2019-5023).

---

**Figure S1.**
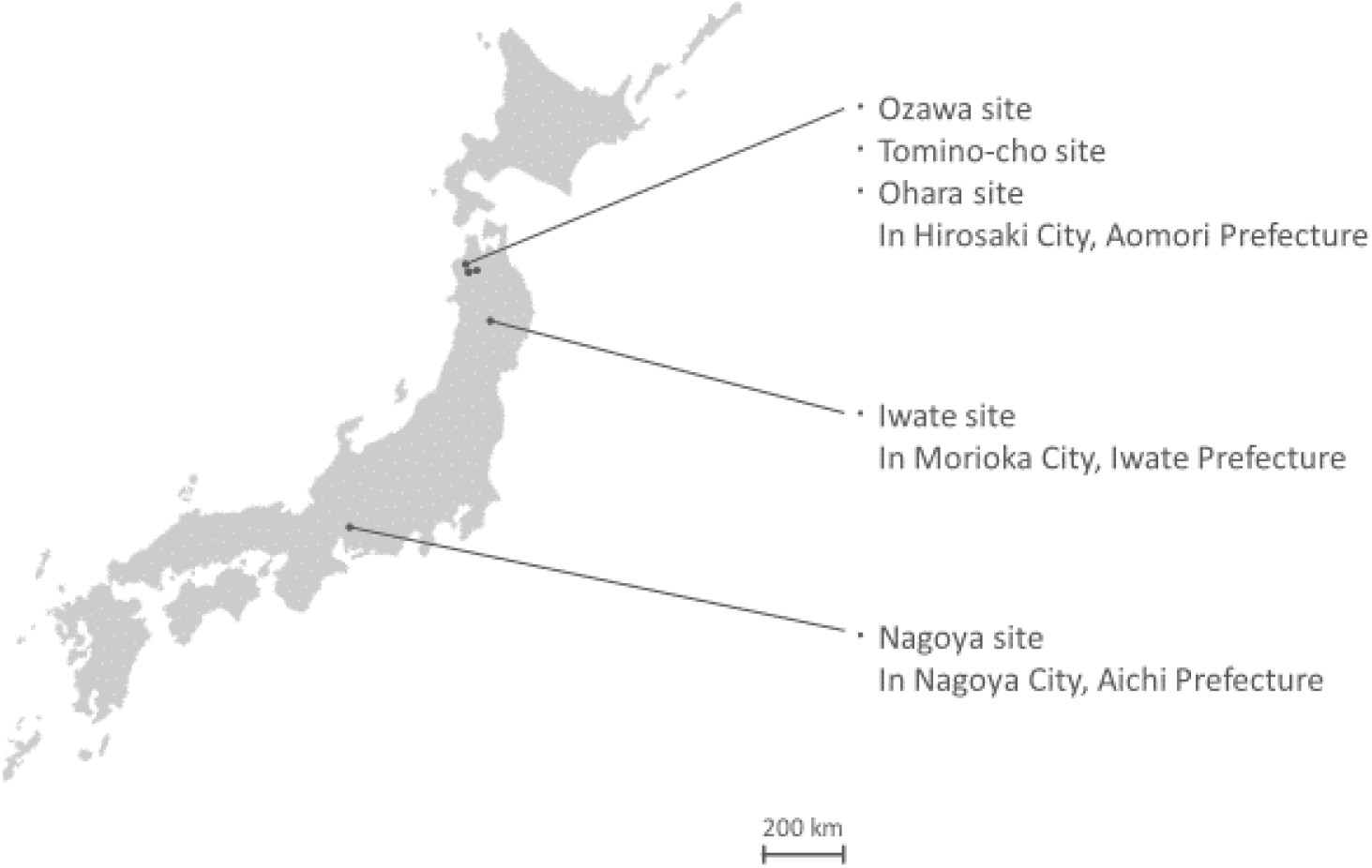
Geographic locations of the five study sites used in the field survey in Japan. These sites were at least 2 km apart.

**Figure S2.**
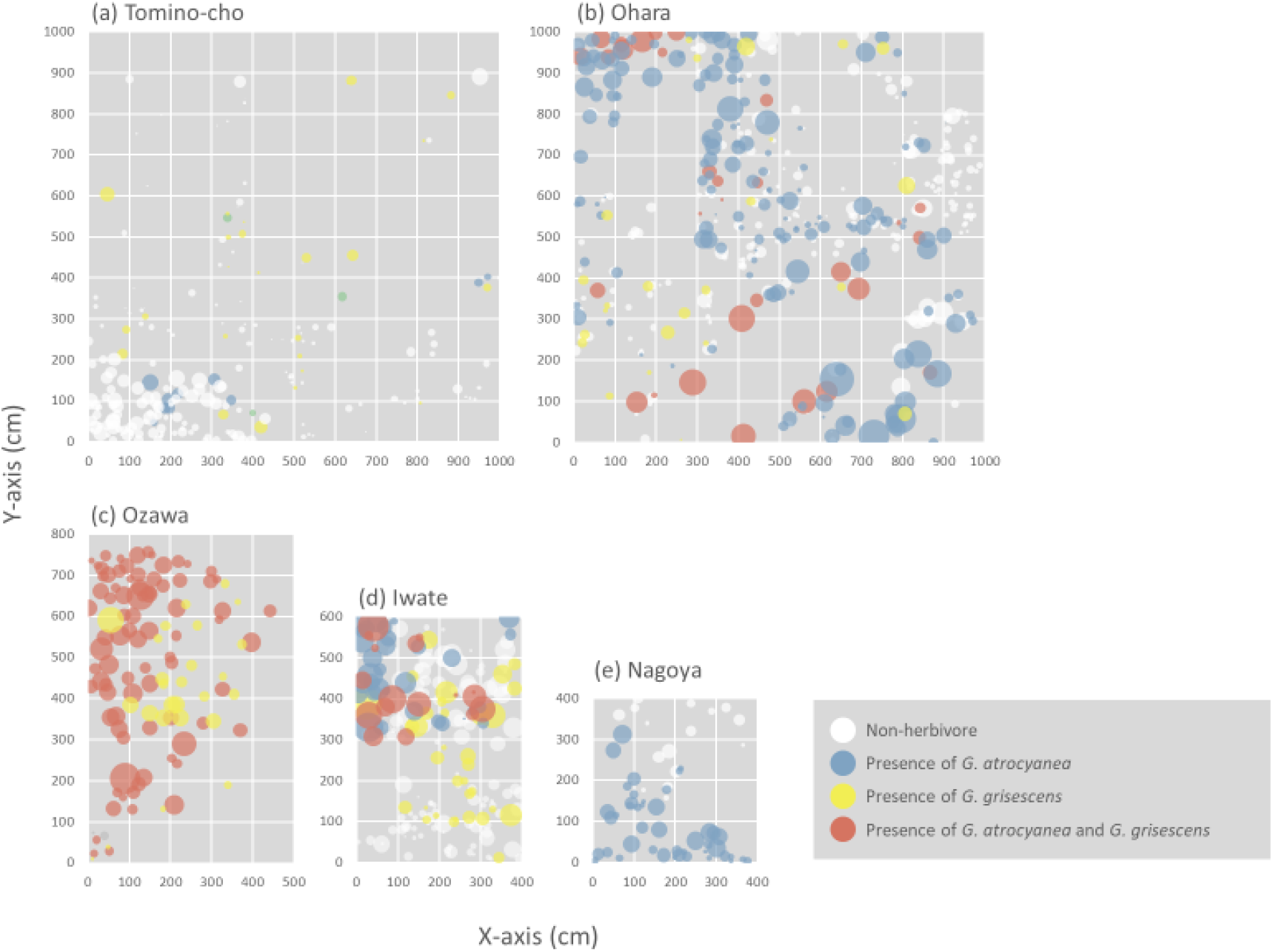
Distributions of *Rumex obtusifolius*, the specialist leaf beetle *Gastrophysa atrocyanea* and the generalist leaf beetle *Galerucella grisescens* at the five study sites. Bubble size in the graphs represents the rosette size of each *R. obtusifolius* plant.

**Figure S3.**
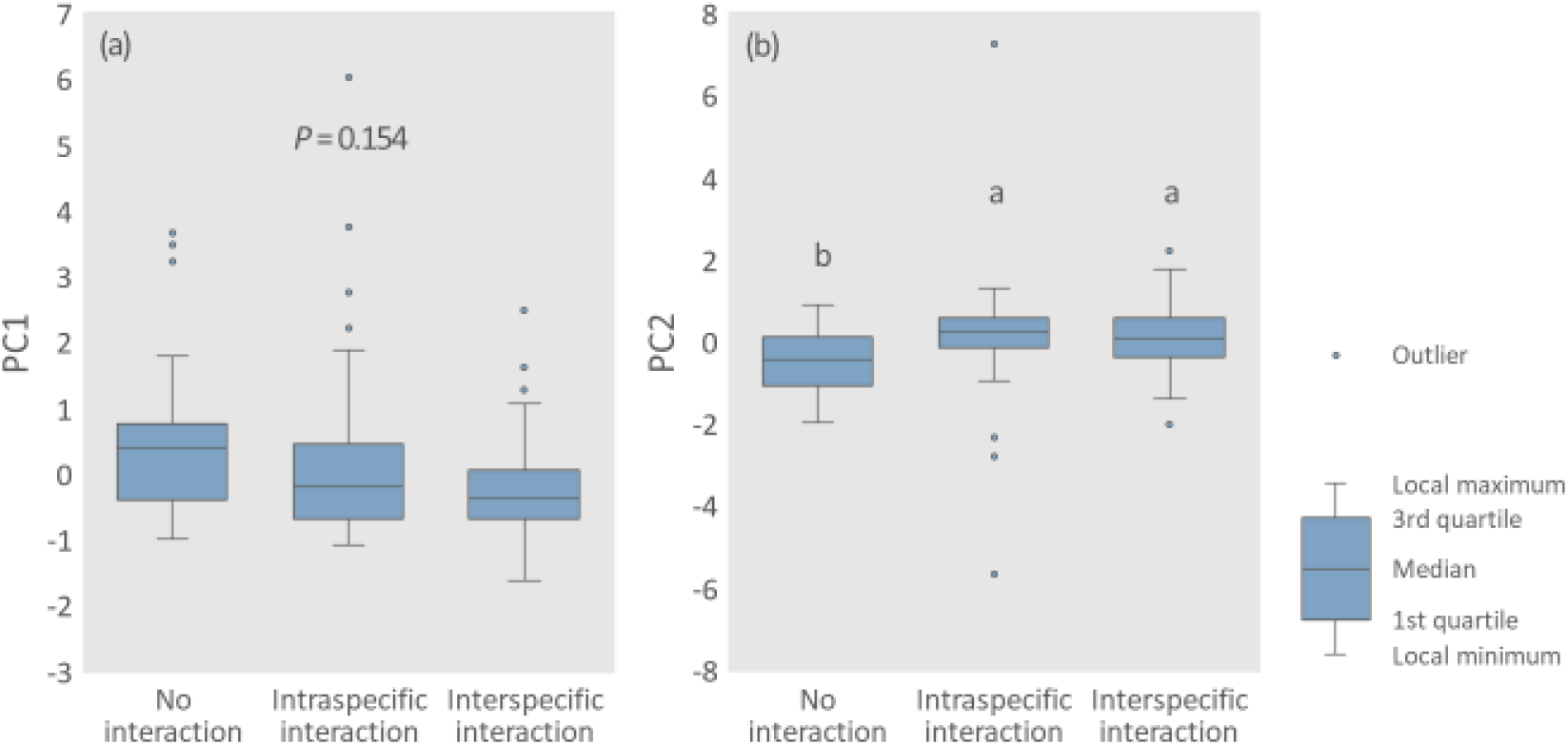
Boxplots of principal component values. (a) PC1 and (b) PC2 for organic acids in the cultivation experiment. Different letters indicate significant differences (GLMM, *P* < 0.05).

## SUPPLEMENTARY METHODS

### Study species

*Rumex obtusifolius* is a perennial herb native to Europe. In the early 1900s, *R. obtusifolius* plants were brought to Japan as contaminants of grasses imported from Europe (Makuchi & Sakai, 1984), and they invaded all regions of Japan except Okinawa. This species grows in wet grassland in Japan. The plants often aggregate with conspecific individuals, and they form patches of various sizes within the population (Van Evert et al., 2011) because their seed germination is promoted by the presence of the leaves of conspecifics via water-soluble-chemical exposure (Ohsaki et al., 2020). The species contains condensed tannins and phenolics as secondary metabolites in the leaves (Feduraev et al., 2019).

*Rumex obtusifolius* leaves are consumed mainly by two leaf beetles (Table 1), *Gastrophysa atrocyane* and *Galerucella grisescen*. *Gastrophysa atrocyanea* is a univoltine species and specialist herbivore of Rumex plants. It occurs on Rumex plant species from spring (March) to mid-summer (July) in Japan (Suzuki, 1985). *Galerucella grisescens* is a multivoltine species and a generalist herbivore of plants of the Polygonaceae family. It occurs on *Rumex* species from spring (March) to early winter (December) in Japan (Suzuki, 1985). Previous studies have shown that larger patches of Rumex plants are associated with *G. atrocyanea* and smaller patches with *G. grisescens* (Suzuki, 1985). This is related to differences in resource utilisation by the two species and to competition advantage: *G. atrocyanea* requires greater amounts of food resources than *G. grisescens*, and G. grisescens, which is less competitive, avoids competition with *G. atrocyanea* (Suzuki, 1985).

### Field survey

#### Total phenolics and condensed tannins analysis

We dried focal plants at 50 °C in an oven over 3 days. Dried leaf tissues were powdered in a mill. Total phenolics were extracted from 20 mg of leaf powder in 10 mL of 50% methanol for 1 h in an ultrasound bath at 40 °C. The content of total phenolics (mg/g) was measured by using the Folin–Ciocalteu method (Julkunen-Tiitto, 1985). Condensed tannins were extracted from 50 mg of dry leaves and were quantified by radial diffusion assay with tannic acid as a standard (Hagerman, 1987).

#### Leaf beetle choice experiment using leaf sections from naturally growing plants

In April 2018, Solitary and Aggregated *Rumex obtusifolius* plants (85 individuals each) with rosette diameters of about 30 cm were selected at random in Hirosaki City. We collected the youngest fully expanded leaves from the plants. These leaves had no damage. We cut one 2-cm piece from the base of each collected leaf. A wet filter paper (8 cm in diameter) was placed in a covered Petri dish (8.5 cm in diameter), and a piece of leaf from a Solitary plant and a piece of leaf from an Aggregated plant were placed on it. Adults of *G. atrocyanea* (36 females and 14 males) and *G. grisescens* (22 females and 13 males) were also collected from the field in Hirosaki City. One beetle was placed in the centre of the Petri dish, which was then kept in a growth chamber for 24 h at 25 °C, with a 12L 12D cycle. Each leaf piece was scanned by using an image scanner, and the consumed area of the leaf was measured by using ImageJ bundled with 64-bit Java 1.8.0_172 image analysis software (Abràmoff et al., 2004). This experimental design is generally adopted for choice experiments with herbivores (e.g., Blüthgen, 2007; Sato et al., 2014; Shirahama et al., 2017; Lackner et al., 2019).

### Cultivation experiments

#### Leaf beetle choice experiment

To reveal whether changes in leaf chemical contents induced by interaction in *R. obtusifolius* influenced the preferences of leaf beetles, we conducted choice experiments with the *R. obtusifolius* leaves used in the cultivation experiment. The experimental design was similar to that described for the choice experiment using field leaves. Leaf pairs were cut into 1.5-cm pieces taken from the leaf base and placed on a Petri dish. The combinations of leaf pairs were as follows: no-interaction treatment versus intraspecific interaction; interspecific interaction versus intraspecific interaction; and no-interaction versus interspecific treatment. In the interspecific interaction treatment, treatments using three different species of plants *(P. asiatica, T. repens* and *F. ovina*) were used equally. In this experiment, we used only *G. atrocyanea*, because in the choice experiment using field leaves only *G. atrocyanea* had expressed a significant preference for group leaves (see Results). Male (*N* = 65) and female (*N* = 69) beetles were collected from Hirosaki City and assigned to the three different interaction treatments (intraspecific interaction versus interspecific interaction, 25 males and 25 females; interspecific interaction versus no interaction, 15 males and 19 females; no interaction versus intraspecific interaction, 25 males and 25 females). Experimental conditions were the same as those in the choice experiment using field leaves.

#### Measurement of leaf traits in cultivated plants

Chlorophyll content. First, a non-destructive chlorophyll meter (SPAD-502 Plus; Konica Minolta, Tokyo, Japan) was used to measure the chlorophyll content in the most recently fully expanded leaves of each cultivated *R. obtusifolius* on the final day of cultivation. A chlorophyll meter is one of the most commonly used diagnostic tools for rapid and non-destructive estimation of chlorophyll content in leaves; the resulting SPAD values are positively correlated with chlorophyll content (Shibaeva et al., 2020). Each leaf was measured twice—in the central part on both sides of the main vein—and the average value was determined. Next, one longitudinal half of each leaf was used to measure primary metabolites. The other halves of the leaves were used in the leaf beetle choice experiment.

#### Organic acids analysis

Organic acids were extracted according to the method of Miyagi et al. (2010). The leaf halves were frozen with liquid nitrogen and stored in a freezer at –80 °C until measurement. Organic acids were extracted according to the method of Miyagi et al. (2010). Frozen leaves (about 50 mg) were milled, and 50% methanol containing 50 mM PIPES (1,4-piperazineethanesulfonate) as an internal standard were added. After initial centrifugation (22,000*g*, 5 min, 4 °C), the supernatant was transferred to a 3-kDa cut-off filter (Millipore, Billerica, MA, USA) and recentrifuged (14,000*g*, 30 min, 4 °C). Five primary metabolites—oxalate, isocitrate, citrate, glycolate and threonate—were selected for analysis. These metabolites had been found in *R. obtusifolius* in our preliminary survey. The resulting filtrate was quantified by capillary electrophoresis triple quadrupole mass spectrometry (CE-QQQ-MS, CE; 7100, MS; 6420 Triple Quad LC/MS, Agilent Technologies, Santa Clara, CA, USA) with multi-reaction monitoring mode according to the method of Miyagi et al. (2019). Because of the small leaf size, plants for which we did not have enough samples for analysis were excluded from the analysis, and the final number of plants analysed for organic acids was slightly smaller than the original collection (no interaction, *N* = 35; intraspecific interaction, *N* = 63; interspecific interaction, *N* = 97).

### Mesocosm experiment

#### Preparation of plants and insects

In August 2018, seeds of *R. obtusifolius* were collected from three individuals in Hirosaki City. Individuals were separated by at least 2 km. In October 2019 and June 2020, seeds were germinated in the same way as described in the cultivation experiment and used for the mesocosm experiments. Two weeks later, we planted two *R. obtusifolius* seedlings in each pot (10.5 cm diameter × 9 cm high) containing seed-free garden soil (Mori Sangyo Co., Ltd, Japan). As a no-interaction treatment, pots were divided in half by a plastic plate to block below-ground interaction of the plants in each experiment (2019, *N* = 35; 2020, *N* = 84), because changes in leaf chemical content in response to conspecific neighbours depend on direct interaction below the ground (Ohsaki, 2020). The other pots were assigned to the interaction treatment, which consisted of two *R. obtusifolius* seedlings with no plastic plate (2019, *N* = 35; 2020, *N* = 74). All pots were watered once a day and maintained at 25 °C, 12L 12D for 30 days in growth chambers. *Gastrophysa atrocyanea* were collected from the field and kept in the laboratory until required for the experiment.

## REFERENCES

Abrahamson, W. G. (2008). Specialization, speciation, and radiation: the evolutionary biology of herbivorous insects. Univ of California Press, Berkeley, CA.

Adler, P. B., Smull, D., Beard, K. H., Choi, R. T., Furniss, T., Kulmatiski, A., Meiners, J. M., Tredennick. A. T., & Veblen, K. E. (2018). Competition and coexistence in plant communities: intraspecific competition is stronger than interspecific competition. Ecology letters, 21, 1319–1329. https://doi.org/10.1111/ele.13098

Akula, R., & Ravishankar, G. A. (2011). Influence of abiotic stress signals on secondary metabolites in plants. Plant Signaling & Behavior, 6, 1720–1731. https://doi.org/10.4161/psb.6.11.17613

Barton, K. E., & Bowers, M. D. (2006). Neighbor species differentially alter resistance phenotypes in *Plantago*. Oecologia, 150, 442–452. https://doi.org/10.1007/s00442-006-0531-z

Broz, A. K., Broeckling, C. D., De-la-Peña, C., Lewis, M. R., Greene, E., Callaway, R. M., Sumner, L. W., & Vivanco. J. M. (2010). Plant neighbor identity influences plant biochemistry and physiology related to defense. BMC Plant Biology, 10, 1–14. https://doi.org/10.1186/1471-2229-10-115

Coutinho, R. D., Cuevas-Reyes, P., Fernandes, G. W., & Fagundes, M. (2019). Community structure of gall-inducing insects associated with a tropical shrub: regional, local and individual patterns. Tropical Ecology, 60, 74–82. https://doi.org/10.1007/s42965-019-00010-7

Craine, J. M., & Dybzinski, R. (2013). Mechanisms of plant competition for nutrients, water and light. Functional Ecology, 27, 833–840. https://doi.org/10.1111/1365-2435.12081

Dudt, J. F., & Shure, D. J. (1994). The influence of light and nutrients on foliar phenolics and insect herbivory. Ecology, 75, 86–98. https://doi.org/10.2307/1939385

Fagundes, M., Barbosa, E. M., Oliveira, J. B., Brito, B. G., Freitas, K. T., Freitas, K. F., & Reis-Junior, R. (2019). Galling inducing Insects associated with a tropical shrub: the role of resource concentration and species interactions. Ecología Austral, 29, 012–019. https://doi.org/10.25260/EA.19.29.1.0.751

Feduraev, P., Chupakhina, G., Maslennikov, P., Tacenko, N., & Skrypnik, L. (2019). Variation in phenolic compounds content and antioxidant activity of different plant organs from *Rumex crispus* L. and *Rumex obtusifolius* L. at different growth stages. Antioxidants, 8, 237. https://doi.org/10.3390/antiox8070237

Feeny, P. (1970). Seasonal changes in oak leaf tannins and nutrients as a cause of spring feeding by winter moth caterpillars. Ecology, 51, 565–581. https://doi.org/10.2307/1934037

Fretwell, D. S., & Lucas, H. L. J. (1969). On territorial behavior and other factors influencing habitat distribution in birds. Acta Biotheoretica, 19, 16–36. https://doi.org/10.1007/BF01601

Godinho, D. P., Janssen, A., Li, D., Cruz, C., & Magalhães, S. (2020). The distribution of herbivores between leaves matches their performance only in the absence of competitors. Ecology and Evolution, 10, 8405–8415. https://doi.org/10.1002/ece3.6547

Goodey, N. A., Florance, H. V., Smirnoff, N., & Hodgson, D. J. (2015). Aphids pick their poison: selective sequestration of plant chemicals affects host plant use in a specialist herbivore. Journal of Chemical Ecology, 41, 956–964. https://doi.org/10.1007/s10886-015-0634-2

Hambäck, P. A., & Beckerman, A. P. (2003). Herbivory and plant resource competition: a review of two interacting interactions. Oikos, 101, 26–37. https://doi.org/10.1034/j.1600-0706.2003.12568.x

Jeschke, V., Kearney, E. E., Schramm, K., Kunert, G., Shekhov, A., Gershenzon, J., & Vassão, D. G. (2017). How glucosinolates affect generalist lepidopteran larvae: growth, development and glucosinolate metabolism. Frontiers in Plant Science, 8, 1995. https://doi.org/10.3389/fpls.2017.01995

Kigathi, R. N., Weisser, W. W., Veit, D., Gershenzon, J., & Unsicker, S. B. (2013). Plants suppress their emission of volatiles when growing with conspecifics. Journal of Chemical Ecology, 39, 537–545. https://doi.org/10.1007/s10886-013-0275-2

Kitazaki, K., Fukushima, A., Nakabayashi, R., Okazaki, Y., Kobayashi, M., Mori, T., Nishizawa, T., Reyes-Chin-Wo, S., Michelmore, R. W., Shoji, K., & Kusano, M. (2018). Metabolic reprogramming in leaf lettuce grown under different light quality and intensity conditions using narrow-band LEDs. Scientific Reports, 8, 1–12. https://doi.org/10.1038/s41598-018-25686-0

Macel, M. (2011). Attract and deter: a dual role for pyrrolizidine alkaloids in plant– insect interactions. Phytochemistry Reviews, 10, 75–82. https://doi.org/10.1007/s11101-010-9181-1

Makuchi, T., & Sakai, H. (1984). Seedling survival and flowering of Rumex obtusifolius L. in various habitats. Weed Research, 29, 123–130. https://doi.org/10.3719/weed.29.123

Mraja, A., Unsicker, S. B., Reichelt, M., Gershenzon, J., & Roscher, C. (2011). Plant community diversity influences allocation to direct chemical defence in *Plantago lanceolata*. PLoS One, 6, e28055. https://doi.org/10.1371/journal.pone.0028055

Muiruri, E. W., Barantal, S., Iason, G. R., Salminen, J. P., Perez-Fernandez, E., & Koricheva, J. (2019). Forest diversity effects on insect herbivores: do leaf traits matter?. New Phytologist, 221, 2250–2260. https://doi.org/10.1111/nph.15558

Nerlekar, A. N. (2018). Seasonally dependent relationship between insect herbivores and host plant density in Jatropha nana, a tropical perennial herb. Biology Open 7, 1–7. https://doi.org/10.1242/bio.035071

Offor, E. (2010). The nutritional requirements of phytophagous insects: why do insects feed on plants? Available at SSRN, 1535274.

Ohsaki, H., Mukai, H., & Yamowo, A. (2020). Biochemical recognition in seeds: Germination of *Rumex obtusifolius* is promoted by leaves of facilitative adult conspecifics. Plant Species Biology, 35, 233–242. https://doi.org/10.1111/1442-1984.12275

Ômura, H. (2018). Plant secondary metabolites in host selection of butterfly. Chemical Ecology of Insects (ed. by J. Tabata), pp. 3–27. CRC Press, Boca Raton, Florida.

Ormeño, E., Bousquet-Mélou, A., Mévy, J. P., Greff, S., Robles, C., Bonin, G., & Fernandez, C. (2007). Effect of intraspecific competition and substrate type on terpene emissions from some Mediterranean plant species. Journal of Chemical Ecology, 33, 277–286. https://doi.org/10.1007/s10886-006-9219-4

Otway, S. J., Hector, A., & Lawton, J. H. (2005). Resource dilution effects on specialist insect herbivores in a grassland biodiversity experiment. Journal of Animal Ecology, 74, 234–240. https://doi.org/10.1111/j.1365-2656.2005.00913.x

R Development Core Team. (2019). R: A language and environment for statistical computing. Vienna, Austria: R Foundation for Statistical Computing.

Rhainds, M., & English-Loeb, G. (2003). Testing the resource concentration hypothesis with tarnished plant bug on strawberry: density of hosts and patch size influence the interaction between abundance of nymphs and incidence of damage. Ecological Entomology, 28, 348–358. https://doi.org/10.1046/j.1365-2311.2003.00508.x

Root, R. B. (1973). Organization of a plant-arthropod association in simple and diverse habitats: the fauna of collards (Brassica oleracea). Ecological Monographs, 43, 95–124. https://doi.org/10.2307/1942161

Scheirs, J., & De Bruyn, L. (2004). Excess of nutrients results in plant stress and decreased grass miner performance. Entomologia experimentalis et applicata, 113, 109–116. https://doi.org/10.1111/j.0013-8703.2004.00215.x

Scherber, C., Eisenhauer, N., Weisser, W.W., Schmid, B., Voigt, W., Fischer, M. et al. (2010). Bottom-up effects of plant diversity on multitrophic interactions in a biodiversity experiment. Nature, 468, 553–556. https://doi.org/10.1038/nature09492

Schoonhoven, L.M., van Loon, J.J.A. & Dicke, M. (2005). Insect–Plant Biology, 2nd edn. Oxford University Press, Oxford.

Sousa-Souto, L., Bocchiglieri, A., Dias, D. D. M., Ferreira, A. S., & José Filho, P. D. L. (2018). Changes in leaf chlorophyll content associated with flowering and its role in the diversity of phytophagous insects in a tree species from a semiarid Caatinga. PeerJ, 6, e5059. https://doi.org/10.7717/peerj.5059

Stephens, A. E., & Myers, J. H. (2012). Resource concentration by insects and implications for plant populations. Journal of Ecology, 100, 923–931. https://doi.org/10.1111/j.1365-2656.2005.00913.x

Suzuki, N. (1985). Resource utilization of three chrysomelid beetles feeding on Rumex plants with diverse vegetational background. Japanese Journal of Ecology, 35, 225–234. https://doi.org/10.18960/seitai.35.2_225

Suzuki, N. (1986). Interspecific competition and coexistence of the two chrysomelids, *Gastrophysa atrocyanea* Motschulsky and *Galerucella vittaticollis* Baly (Coleoptera: Chrysomelidae), under limited food resource conditions. Ecological Research, 1, 259–268. https://doi.org/10.1007/BF02348683

Takigahira, H., & Yamawo, A. (2019). Competitive responses based on kin-discrimination underlie variations in leaf functional traits in Japanese beech (*Fagus crenata*) seedlings. Evolutionary Ecology, 33, 521–531. https://doi.org/10.1007/s10682-019-09990-3

Titayavan, M., & Altieri, M. A. (1990). Synomone-mediated interactions between the parasitoid *Diaeretiella rapae* and *Brevicoryne brassicae* under field conditions. Entomophaga, 35, 499–507. https://doi.org/10.1007/BF02375084

Tuller, J., Queiroz, A. C. M., Luz, G. R., & Silva, J. O. (2013). Gall-forming insect attack patterns: a test of the Plant Vigor and the Resource Concentration Hypotheses. Biotemas, 26, 45–51. https://doi.org/10.5007/2175-7925.2013v26n1p45

War, A. R., Paulraj, M. G., Ahmad, T., Buhroo, A. A., Hussain, B., Ignacimuthu, S., & Sharma, H. C. (2012). Mechanisms of plant defense against insect herbivores. Plant Signaling & Behavior, 7, 1306–1320. https://doi.org/10.4161/psb.21663

Wheat, C. W., Vogel, H., Wittstock, U., Braby, M. F., Underwood, D., & Mitchell-Olds, T. (2007). The genetic basis of a plant–insect coevolutionary key innovation. Proceedings of the National Academy of Sciences, 104, 20427–20431. https://doi.org/10.1073/pnas.0706229104

Yamamura, K. (1999). Relation between plant density and arthropod density in cabbage fields. Population Ecology, 41, 177–182. https://doi.org/10.1007/s101440050020

Yamawo A. (2015) Relatedness of neighboring plants alters the expression of indirect defense traits in an extrafloral nectary-bearing plant. Evolutionary Biology, 42, 12–19. https://doi.org/10.1007/s11692-014-9295-2

Yamawo A. (2021). Intraspecific competition favors ant-plant protective mutualism. Plant Species Biology. https://doi.org/10.1111/1442-1984.12331

Yamawo, A., & Mukai, H. (2020). Outcome of interspecific competition depends on genotype of conspecific neighbours. Oecologia, 193, 415–423. https://doi.org/10.1007/s00442-020-04694-w

## References For Supplementary Methods

Abràmoff, M. D., Magalhães, P. J., & Ram, S. J. (2004). Image processing with ImageJ. Biophotonics international, 11, 36–42.

Blüthgen, N., & Metzner, A. (2007). Contrasting leaf age preferences of specialist and generalist stick insects (Phasmida). Oikos, 116, 1853–1862. https://doi.org/10.1111/j.0030-1299.2007.16037.x

Hagerman, A. E. (1987). Radial diffusion method for determining tannin in plant extracts. Journal of Chemical Ecology, 13, 437–449. https://doi.org/10.1007/BF01880091

Julkunen-Tiitto, R. (1985). Phenolic constituents in the leaves of northern willows: methods for the analysis of certain phenolics. Journal of Agricultural and Food Chemistry, 33, 213–217. https://doi.org/10.1021/jf00062a013

Lackner, S., Lackus, N. D., Paetz, C., Köllner, T. G., & Unsicker, S. B. (2019). Aboveground phytochemical responses to belowground herbivory in poplar trees and the consequence for leaf herbivore preference. Plant, Cell & Environment, 42, 3293–3307. https://doi.org/10.1111/pce.13628

Makuchi, T., & Sakai, H. (1984). Seedling survival and flowering of *Rumex obtusifolius* L. in various habitats. Weed Research, 29, 123–130. https://doi.org/10.3719/weed.29.123

Miyagi, A., Noguchi, K., Tokida, T., Usui, Y., Nakamura, H., Sakai, H., Hasegawa, T., & Kawai-Yamada, M. (2019). Oxalate contents in leaves of two rice cultivars grown at a free-air CO2 enrichment (FACE) site. Plant Production Science, 22, 407–411. https://doi.org/10.1080/1343943X.2019.1598272

Miyagi, A., Takahashi, H., Takahara, K., Hirabayashi, T., Nishimura, Y., Tezuka, T., Kawai-Yamada, M., & Uchimiya, H. (2010). Principal component and hierarchical clustering analysis of metabolites in destructive weeds; polygonaceous plants. Metabolomics, 6, 146–155. https://doi.org/10.1007/s11306-009-0186-y

Ohsaki H. (2020). Effects of plant-plant interactions on resource usage of herbivores. MS thesis, Hirosaki University, Hirosaki (in Japanese)

Sato, Y., Kawagoe, T., Sawada, Y., Hirai, M. Y., & Kudoh, H. (2014). Frequency-dependent herbivory by a leaf beetle, *Phaedon brassicae*, on hairy and glabrous plants of *Arabidopsis halleri* subsp. gemmifera. Evolutionary Ecology, 28, 545–559. https://doi.org/10.1007/s10682-013-9686-3

Shibaeva, T. G., Mamaev, A. V., & Sherudilo, E. G. (2020). Evaluation of a SPAD-502 Plus Chlorophyll Meter to Estimate Chlorophyll Content in Leaves with Interveinal Chlorosis. Russian Journal of Plant Physiology, 67, 690–696. https://doi.org/10.1134/S1021443720040160

Shirahama, S., Yamawo, A., & Tokuda, M. (2017). Dimorphism in trichome production of *Persicaria lapathifolia* var. *lapathifolia* and Its multiple effects on a leaf beetle. Arthropod-Plant Interactions, 11, 683–690. https://doi.org/10.1007/s11829-017-9520-x

Suzuki, N. (1985). Resource utilization of three chrysomelid beetles feeding on *Rumex* plants with diverse vegetational background. Japanese Journal of Ecology, 35, 225–234. https://doi.org/10.18960/seitai.35.2_225

Van Evert, F. K., Samsom, J., Polder, G., Vijn, M., Dooren, H. J. V., Lamaker, A., van der Heijden, G. W., Kempenaar, C., van der Zalm, T., & Lotz, L. A. P. (2011). A robot to detect and control broad-leaved dock (Rumex obtusifolius L.) in grassland. Journal of Field Robotics, 28, 264–277. https://doi.org/10.1002/rob.20377

